# Integrating genetic and gene expression data in network-based stratification analysis of cancers

**DOI:** 10.1101/2024.10.18.619017

**Authors:** Kenny Liou, Ji-Ping Wang

## Abstract

Cancers are complex diseases that have heterogeneous genetic drivers and varying clinical outcomes. A critical area of cancer research is organizing patient cohorts into subtypes and associating subtypes with clinical and biological outcomes for more effective prognosis and treatment. Large-scale studies have collected a plethora of omics data across multiple tumor types. These studies provide an extensive dataset for stratifying patient cohorts. Network-based stratification (NBS) approaches have been presented to classify cancer tumors using somatic mutation data. A challenge in cancer stratification is integrating omics data to yield clinically meaningful subtypes. In this study, we integrate somatic mutation data with RNA sequencing data within the NBS framework and investigate the effectiveness of integrated NBS on three cancers: ovarian, bladder, and uterine cancer. We show that integrated NBS subtypes are more significantly associated with overall survival or histology. Integrated NBS networks also reveal highly influential genes that drive cancer initiation and progression. This comprehensive approach underscores the significance of integrating genomic data types in cancer subtyping, offering profound implications for personalized prognosis and treatment strategies.

## Introduction

Cancer, a multifaceted disease, exhibits remarkable genetic diversity among patients, rendering the development of effective treatments a formidable challenge. To gain valuable insights into this complexity, large-scale initiatives like the International Cancer Genome Consortium (ICGC) [1] and The Cancer Genome Atlas (TCGA) [2] meticulously profile genomic, transcriptomic, and epigenomic data. This includes DNA methylation, microRNA expression, and protein expression data, unveiling a treasure trove of information. Consequently, the massive amount of data from these projects has spurred a demand for informatics solutions to unearth molecular pathways dictating tumor progression, understanding disease populations, and designing precision-targeted treatment strategies [3].

In cancer informatics, a pivotal objective is to categorize or stratify heterogeneous cancer tumors into clinically and biologically meaningful subtypes using molecular profiling data. Historically, this endeavor primarily leveraged mRNA expression data, yielding valuable insights into cancers like ovarian cancer [4] and breast cancer [5]. However, it has fallen short in other cancer types, such as colorectal cancer, where the association between molecular subtypes and clinical phenotypes remains elusive [6]. This limitation may be attributed to issues like sample quality and overfitting, inherent to gene expression analysis [7].

As the landscape of genome-scale data diversifies and expands, there emerges an urgent need to develop methodologies that effectively integrate multi-omics data for more robust tumor subtyping [8]. Multi-omic research in cancer genetics offers promising results, such as in breast cancer [9, 10]. Typically, multi-omic integration methods synthesize data after processing -omics data types separately or using different -omics data types for different parts of the methodology pipeline [8]. The advantage of multi-omics is that the potential interactions between various molecular layers are considered, providing a holistic perspective that can lead to different results compared to single-data-type methods [11]. In this paper, we integrate molecular profiles before processing these data types to explore if pre-process integration can yield meaningful results.

Recent years have witnessed the widespread adoption of gene interaction networks, with cancer being increasingly recognized as a network-driven disease [12]. Somatic mutation profiles harbor cancer-driver genes capable of instigating genetic mutations in other genes. Network-based Stratification (NBS) [13] is an approach that melds gene interaction networks with somatic mutation profiles. This fusion involves mapping somatic mutation profiles onto a cancer network and propagating these mutations throughout the network to create smoothed network profiles. NBS uses these profiles to stratify tumors by clustering patients with similar smoothed network profiles.

In line with these trends in cancer genomics, we propose a multi-omics methodology grounded in NBS for tumor stratification, fusing somatic mutation profiles with RNA gene expression data. We fuse somatic mutation and gene expression profiles before applying network propagation to the integrated profiles to explore if integrating profiles before network smoothing can generate informative cancer subtypes. By integrating these two data types before network smoothing, we have successfully generated robust, biologically significant, and clinically informative tumor subtype clusters. Our methodology was tested using the TCGA ovarian, uterine, and bladder carcinoma cohorts, yielding compelling results that can shed new light on understanding cancer population structures through an integrated approach.

## Materials and methods

### Integrating genetic profiles

Ovarian serous adenocarcinoma, uterine endometrial carcinoma, and bladder urothelial carcinoma somatic mutation and gene expression data were downloaded from the TCGA data portal in June 2023. Only individuals with somatic mutation and gene expression data were retained for the experiment. 279 ovarian cancer, 318 uterine endometrial carcinoma, and 399 bladder urothelial carcinoma patients from the TCGA cohorts were used in subsequent analysis. Somatic mutation profiles are binary vectors where a ’0’ or ’1’ for whether a gene for that individual has no mutation or a mutation, respectively. Gene expression profiles are continuous TPM normalized values where the value represents the level of gene expression. The gene expression profiles were min-max normalized gene by gene to match the 0 to 1 range of somatic profiles. Somatic mutation and gene expression profile integration can be represented through this formula:

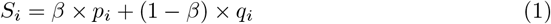

where 0 *< β <* 1 is a tuned hyperparameter chosen by the user to linearly combine the somatic mutation profile *p*_*i*_ and the normalized gene expression profile *q*_*i*_ to result in the integrated profile *S*_*i*_ for individual *i*. In the integration process of NBS for ovarian cancer, bladder cancer, and uterine cancer, we utilized tuned *β* values of 0.8, 0.3, and 0.1, respectively. The value of *β* for ovarian and bladder cancers were chosen to give the most significant p-value in the log-rank test from Kaplan Meier survival analysis or the log-likelihood ratio test from Cox regression survival analysis. Survival analysis for uterine cancer was impeded by the notably low mortality rate within the cohort, alongside the absence of significant associations with survival observed among both single-data type and integrated subtypes. To assess the association of uterine cancer subtypes, we examined the association of TCGA subtypes to our generated Multi-NBS subtypes (*χ*^2^ association test statistic) to derive the value of *β*. Additional details about TCGA are provided at www.cancer.gov/ccg/research/genome-sequencing/tcga.

### Gene interaction network

The constructed network is derived from PCNet [16], a network with 19,781 genes and 2,724,724 interactions that is then filtered for cancer-specific genes and interactions in at least one of these four sources: [17–20]. The resulting cancer subnetwork contained 2291 nodes. This network was used for all three methods of NBS. Further details on constructing this network can be found in a previous NBS publication [15].

### Network propagation

The integrated profiles are mapped onto the gene interaction network and then network propagation [44] is applied to diffuse the signals across the network. Let *m* be the number of genes, *n* be the number of patients, *F*_0_ be the initial *n × m* (patient *×* gene matrix), and *A* be the symmetric adjacency matrix (*m × m*) representing the gene-gene interaction network obtained above. Network propagation follows the following iterative procedure:

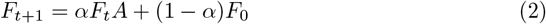

where *α* = 0.7, which is derived from benchmarking results reported in earlier NBS publications [13]. The network is propagated till *F*_*t*_ converges (|*F*_*t*+1_ − *F*_*t*_ *<* 0.001|). After convergence, the resulting matrix *F*_*t*_ was quantile normalized by row (patient) to ensure each patient followed the same distribution. *F* represents the final normalized and smoothed integrated matrix.

### Network-regularized NMF

Non-negative matrix factorization (NMF) decomposes a matrix into two non-negative matrices whose product results in the original matrix. Network-regularized NMF is an extension of NMF that constrains NMF to respect the network structure [45–47]. The following objective function is minimized:

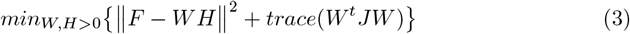

*F* is approximately decomposed into the product of non-negative matrices *W* (*m* by *K* matrix) and *H* (*K* by *n* matrix). *W* is a collection of basis vectors (“meta-genes”) and *H* represents the basis loadings. *K* controls the dimension reduction and we used values of *K* = 2, 3, 4, 5, 6, 7, 8 in the following context. The *trace*(*W*^*t*^*JW*) function is responsible for constraining the basis vectors in *W* to respect the local neighboring network structure. *J* represents the graph Laplacian of the *k*-nearest neighbor network and we used *k* = 11 as described in previous work [13].

### Consensus clustering

We used consensus clustering [48] to ensure robust clustering to achieve the final patient cluster assignments. We performed network-regulated NMF using a random sampling without replacement of 80% of patients. This was repeated 100 times as described previously [13]. The collection of 100 clustering results was used to construct a similarity matrix that recorded the frequency with which patient pairs had the same cluster assignment from all iterations where both patients in the pair were sampled.

### Implementation of Integrated Network-Based Stratification

The implementation of integrated network-based stratification is a Python 3.10 version based on an existing Python 2.7 implementation of NBS [15].

### Cluster analysis

We used Silhouette scores [21] as internal cluster measures. The Silhouette score calculates how well a patient is affiliated with its cluster compared to neighboring clusters. We also used Adjusted Mutual Information (AMI) [22] to assess cluster similarity between clusters formed through different data-type generated clusters.

### Survival analysis

Survival analysis was performed through the lifelines [49]and scikit-survival package [50]. Kaplan-Meier survival curves [51] were fitted to the subtypes generated and log-rank tests were performed on single-data type and integrated NBS subtypes to assess the association between the subtypes and survival. We also fitted a semi-parametric Cox proportional hazard model [52]. The Cox hazard model can be represented with a hazard function *h*(*t*|*X*_*i*_) at time *t* for an individual *i*, given *p* covariates, denoted by *X*_*i*_:

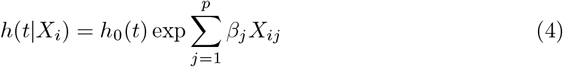

We included covariates such as age, race, and gender in addition to the cluster assignments to assess cluster assignment influence on survival. The log-likelihood ratio test compares the full model with subtype assignments and clinical covariates against a null model. The log-likelihood ratio test and the associated p-value provide an estimate of the predictive power of the full model (clinical covariates and subtype assignments) compared to the null model. The concordance index assesses the discriminatory power of the model by evaluating the ability of the model to correctly rank the survival time of pairs of patients. Maximizing the concordance index indicates the model is discriminative of early events (which are associated with higher-risk patients) and later events.

### Association with TCGA subtypes

Association with TCGA subtypes is calculated through Pearson’s *χ*^2^ test of independence between computed integrated subtypes or single-data type subtypes and documented TCGA subtypes. The documented TCGA subtypes are obtained through the R “TCGAbiolinks” library [23–25], which are determined through genomic, transcriptomic, and proteomic characterization of tumors using array and sequence-based technologies by The Cancer Genome Atlas Research Network [26–28].

### Identifying high-scoring genes

The integrated profiles were first propagated through the network propagation process described above. The propagated integrated profiles are then grouped by the subtype assignments generated using Multi-NBS. After being grouped by subtype the propagated integrated profiles are averaged by number of patients in each subtype to find genes with the highest average hybrid score in the network for each integrated profile subtype. We refer to the score as a ”hybrid score” because the network values for integrated profile networks are a combination of gene expression and somatic mutation data.

For somatic mutation high-scoring genes, the exact same process is applied except we only used somatic mutation profiles and the subtype assignments generated from using somatic mutation profile NBS. We first applied propagation to the somatic mutation profiles. The propagated somatic mutation profiles are grouped by the subtype assignments given by NBS that used only somatic mutation profiles. After being grouped, the propagated somatic mutation profiles are averaged by number of patients in each subtype to find genes with the highest average mutation score in the network for each somatic mutation profile subtype. Note that we refer to the somatic mutation scores as mutation scores because the network values are derived from solely somatic mutation data.

## Results

For the three cancer types under consideration, ovarian, uterine, and bladder carcinoma, only individuals extracted from the TCGA database with both somatic mutation and gene expression profiles were retained, giving 279 ovarian carcinoma, 318 uterine endometrial carcinoma, and 399 bladder urothelial carcinoma patients. We did not remove individuals with fewer than 10 somatic mutations. Multi-omic Network-Based Stratification (Multi-NBS) merges somatic mutation profiles with RNA sequencing (RNA-seq) gene expression data through a linear combination to creating an integrated profile (Fig 1). The resulting integrated matrix is then processed through NBS to generate patient subtype assignments. The underlying network used in this context is a sub-network comprised of cancer-related genes, which was constructed from PCNet [16], but filtered using four databases containing cancer-specific genes and interactions [17–20]. To assess the efficacy of the subtypes generated by Multi-NBS, we conduct a rigorous performance evaluation, comparing them against subtypes derived from single-data type NBS. Additionally, we employ benchmarking procedures to identify the optimal *β* values for integration, ensuring the robustness and informativeness of our results (See Methods).

**Fig 1.**
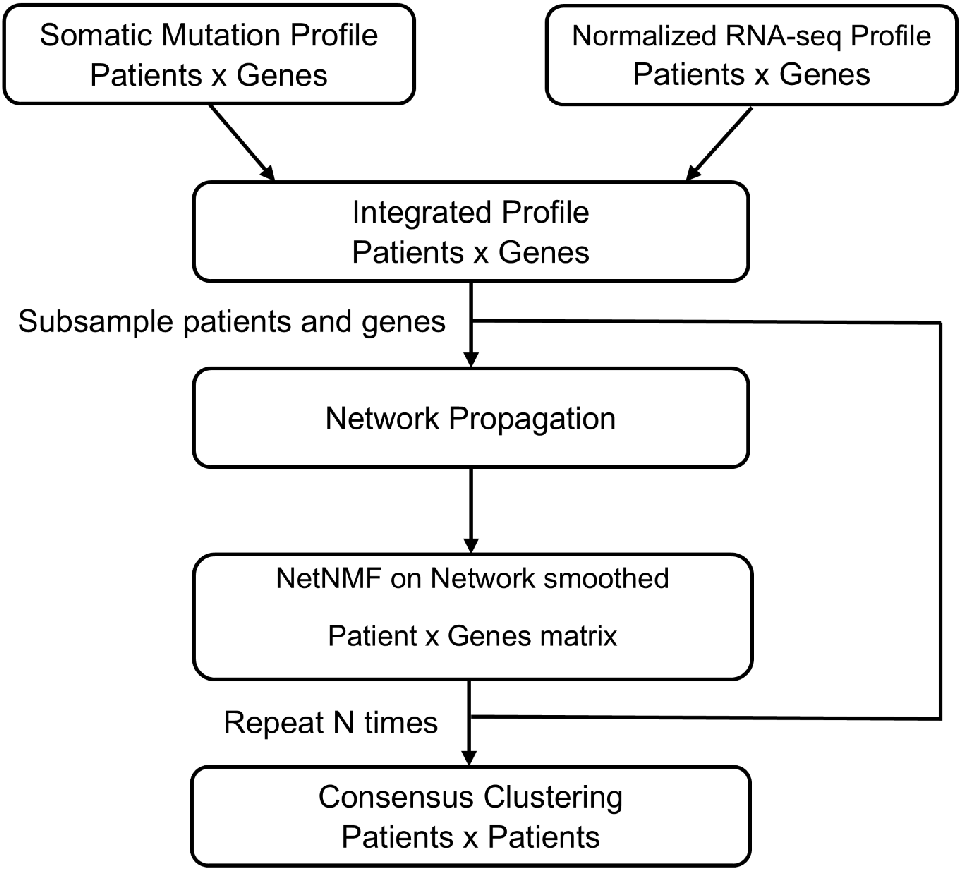
Integrated NBS workflow. A workflow of an integrated approach to network-based stratification.

In the following we assess the efficacy of Multi-NBS clusters in terms of clustering metrics, clinical significance in predicting survival time, and association with histological characteristics by benchmarking with clusters obtained from single data type. In addition we will investigate whether the identified Multi-NBS clusters can reveal new high-scoring genes within the networks to that have cancer relevance.

### Cluster evaluation

We first evaluated whether the clusters generated by Multi-NBS are comparable to single-data-type clusters using Silhouette scores [21]. Higher Silhouette scores (values closer to 1) indicate well-formed clusters while lower scores (values closer to -1) indicate poorly-formed clusters. Values close to 0 in Silhouette scores are still generally considered good classification. We plotted the Silhouette scores as a function of the number of clusters generated under each method (Fig 2). Somatic mutation clusters appeared to have the poorest formed clusters while RNA-seq and Multi-NBS clusters appeared to generate slightly more well-formed clusters (Fig 2A and Fig 2B), while RNA-seq seemed to generate the most well-formed clusters (Fig 2C). The Multi-NBS clusters did not show a uniformly better pattern than other two. Instead, in ovarian and bladder cancers, we see that the Multi-NBS Silhouette score curves were similar to RNA-seq clusters, while in uterine cancer, the Multi-NBS Silhouette score was observed to be between the two single-data-type Silhouette scores.

**Fig 2.**
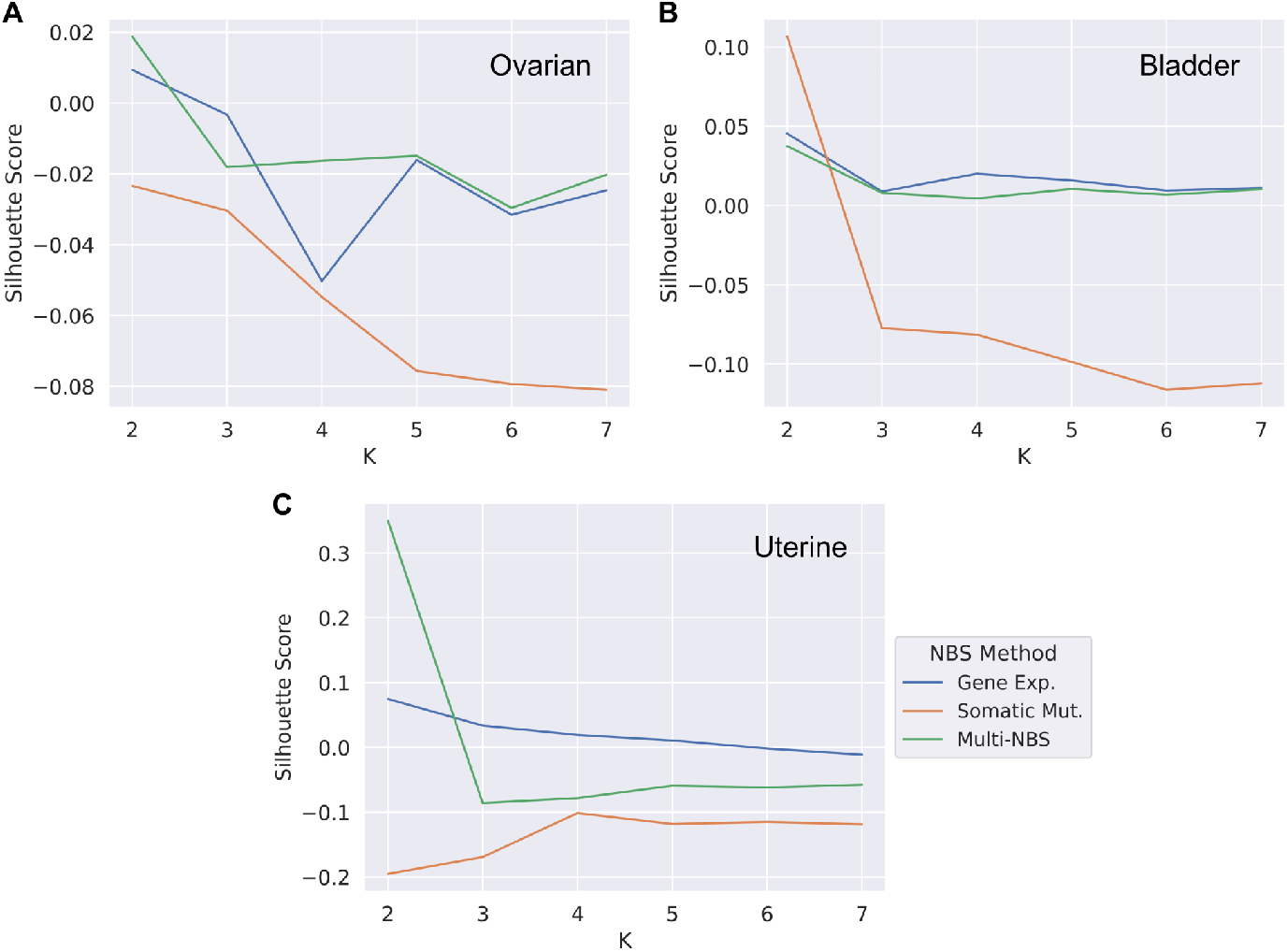
Silhouette score analysis. Silhouette scores of single-data-type generated clusters and Multi-NBS generated clusters. Refer to the legend in subfigure (C) for all subfigures. (A) Silhouette scores of ovarian cancer clusters. (B) Silhouette scores of bladder cancer clusters. (C) Silhouette scores of uterine cancer clusters.

We next compared the similarity of cluster contents using Adjusted Mutual Information (AMI) [22]. AMI is a measure of cluster independence, where a score of 0 means the clusters are completely independent and a score of 1 means the clusters are completely identical. We observed that across all cancer types, somatic mutation and RNA-seq clusters were mostly independent of each other (Fig 3). The Multi-NBS and RNA-seq clusters showed relatively higher cluster similarity and the AMI values are 0.28, 0.82 and 0.8 respectively for ovarian, bladder and uterine cases, reflecting the impact of RNA-seq data in the calculation of blended gene profile *S*_*i*_ (see Eq 1) (i.e., 1 − *β* are 0.2, 0.8 and 0.9 respectively). However, we see the AMI scores between Multi-NBS and somatic mutation clusters are unproportionally lower in all three cases, possibly because the somatic mutation profile is binary while RNA-seq profile is continuous such that the integrated profile more resembles the RNA-seq profile. In the following we shall investigate how the blended gene profile may benefit characterization of the cancer subtypes clinically.

**Fig 3.**
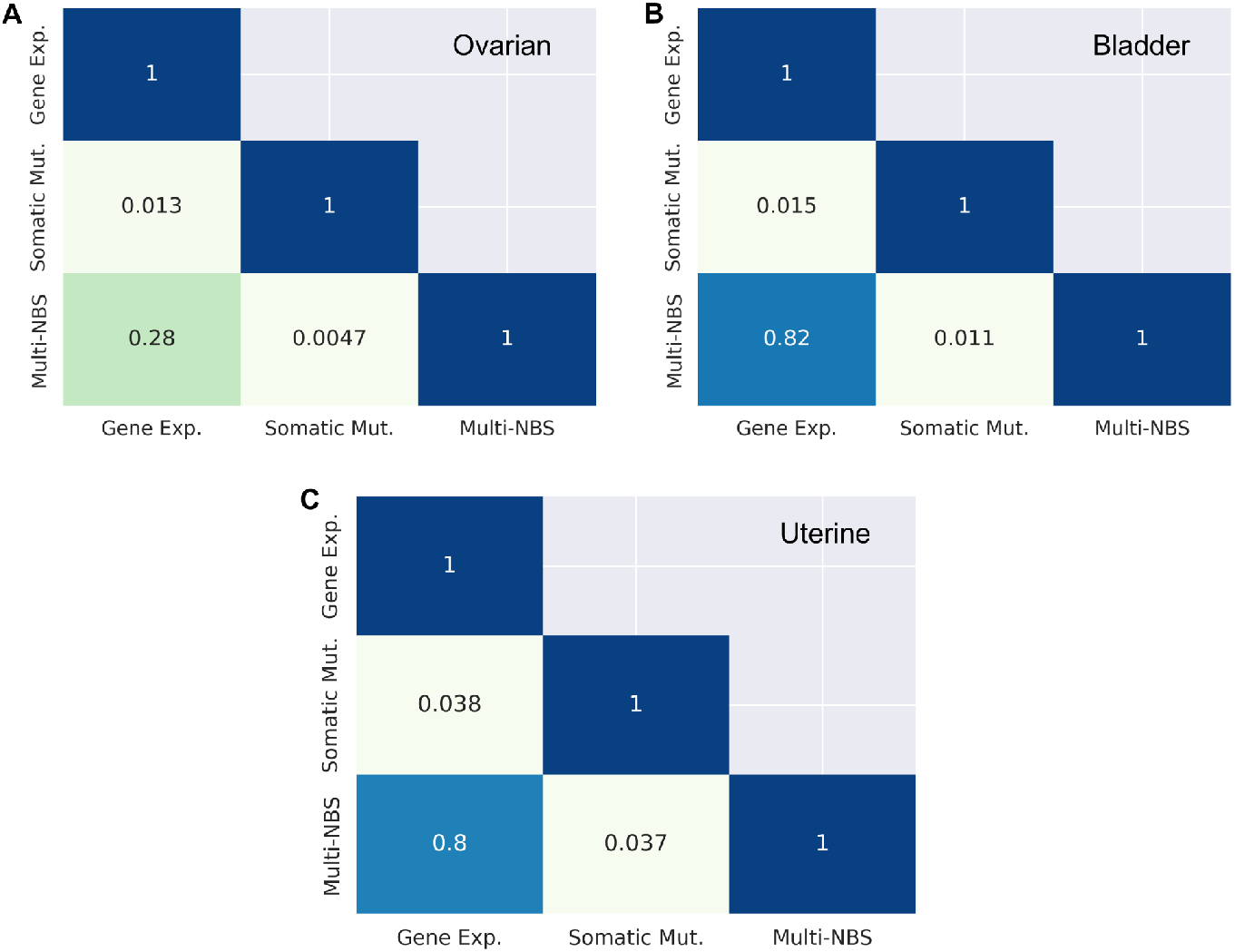
Adjusted mutual information scores. Adjusted Mutual Information scores of single-data-type generated clusters and Multi-NBS generated clusters. (A) AMI scores for ovarian cancer clusters. (B) AMI scores for bladder cancer clusters. (C) AMI scores for uterine cancer clusters.

### Survival analysis

To further assess the generated subtypes, we first examined the Kaplan Meier estimate of the survival function of the generated cancer subtypes, and then a Cox’s proportional hazard regression model to investigate how age, gender, and race as covariates may affect to survival in addition to the cluster assignments. We found that uterine cancer subtypes, whether identified through single-data type NBS or integrated NBS, did not show significant associations with survival due to the low mortality rate within the cohort at significance level *α* = 0.05 (to be used throughout the following context). For instance, uterine cancer subtypes derived solely from somatic mutation data for *K* = 3 did not exhibit a significant association with survival (S3 Fig, log-rank test, *p* = 3.837 *×* 10^−1^). These findings are consistent with previous work [13]. Consequently, further survival analysis for uterine cancer was not pursued.

In ovarian cancer, we first consider *K* = 4 Multi-NBS subtypes, as it is classified into 4 subtypes based on cancer histology. We note that Multi-NBS subtypes were significantly associated with Kaplan Meier survival functions (Fig 4A, log-rank test, *p* = 6 *×* 10^−3^ for *K* = 4). In contrast, subtypes derived from somatic mutation or RNA-seq profiles for *K* = 4 were not found to be significantly associated with survival (S1 Fig, log-rank test, *p* = 1.79 *×* 10^−1^ and *p* = 1.423 *×* 10^−1^, respectively). However, we did observe ovarian cancer subtypes generated from somatic mutation data alone for *K* = 2 are significantly associated with survival, consistent with prior work (S3 Fig, log-rank test, *p* = 9.8 *×* 10^−3^) [13, 14]. The difference in the p-value is mostly likely due to the different samples used in each of these studies.

**Fig 4.**
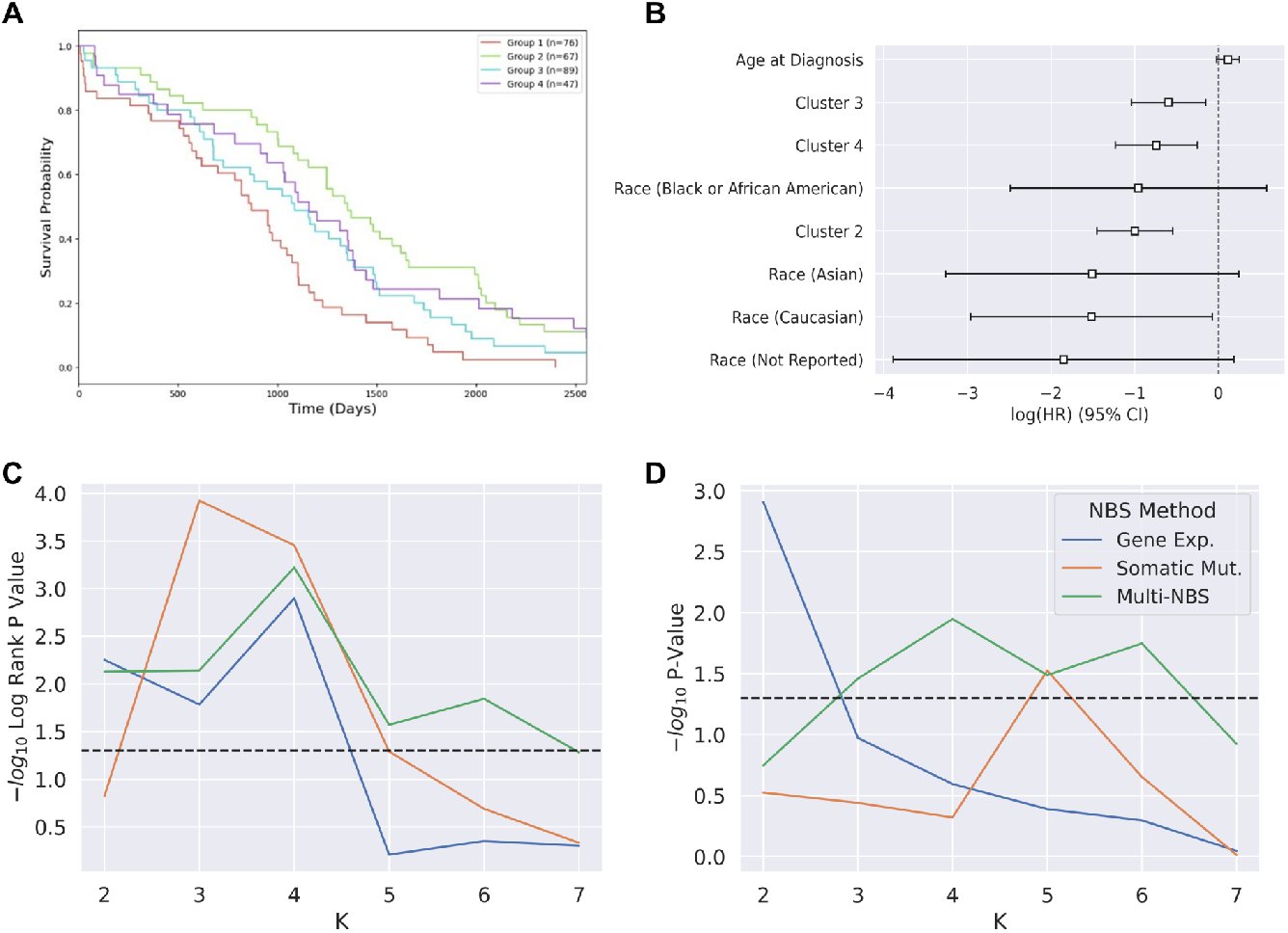
Ovarian cancer survival analysis. Ovarian cancer survival analysis indicate that integrated subtypes are significantly informative of survival. Refer to the legend in subfigure (D) for subfigures (C) and (D). (A) Survival curves indicating the probability of survival for subtypes generated by Multi-NBS (log-rank test, *p* = 6 × 10^−3^). (B) The Cox regression log hazard ratios for *K* = 4 integrated subtypes. (C) Association of NBS ovarian cancer subtypes to survival time through the log-rank test. The black line represents *p* = 0.05. (D) The predictive power of generated OV subtypes on survival through the log-likelihood ratio test. The black line represents *p* = 0.05.

In the Cox-regression model, we found the log hazard ratio for cluster subtypes and Caucasian covariates were all significantly different from zero in Fig 4B. The observed negative log hazard ratio of the three cluster assignments indicated that subtypes 2 through 4 had a significantly lower hazard compared to subtype 1. This result confirmed the Kaplan Meier estimates of the survival function, where group 1 showed the shortest survival time. In contrast for the single-data type clusters, we found the log hazard ratio for the subtypes did not significantly differ from zero (S1 Fig).

When we varied *K* for the subtypes definition, Multi-NBS subtypes exhibited more informative survival patterns across various values of *K*, demonstrating comparable or superior performance in associating subtypes with survival outcomes compared to subtypes derived from single-data types (Fig 4C). Notably, subtypes derived solely from somatic mutation profiles also show significant associations with Kaplan Meier survival functions, consistent with prior work on somatic mutation NBS subtypes [13]. When we accounted for clinical covariates such as age and race, we found Multi-NBS continued to consistently yield robust subtypes that remained more predictive of survival than somatic mutation or RNA-seq profiles for *K >* 2 (Fig 4D).

For bladder cancer, we found that Multi-NBS subtypes proved highly predictive of patient survival time (Fig 5A, log-rank test, *p* = 2 *×* 10^−3^ for *K* = 4). Notably, the most aggressive bladder cancer Multi-NBS subtype exhibited an average survival time of 1100 days (36 months), while the least aggressive bladder cancer subtype showed an average survival time exceeding 2500 days (82 months). We found subtypes generated using only somatic mutation profiles or RNA-seq profiles were not significantly associated with survival (S2 Fig log-rank test, *p* = 1.989 *×* 10^−1^ and *p* = 7.51 *×* 10^−2^ for *K* = 4, respectively). Log hazard ratios for *K* = 4 show that all cluster assignments coefficients except for the subtype 2 coefficient significantly differ from zero (Fig 5B). Similar to ovarian cancer, we see all single-data type cluster assignment coefficients do not significantly differ from zero, indicating that the subtypes generated from mutation or RNA-seq data alone are less informative of survival when covariates are taken into account (S2 Fig). We also see Multi-NBS BLCA subtypes had comparable predictive power to RNA-seq generated subtypes for survival time even after adjusting for clinical covariates such as age, gender, and race in Fig 5C. This highlights the potential utility and clinical relevance of Multi-NBS in delineating patient subtypes with distinct survival outcomes across cancer types, yielding subtypes that may be more informative of survival compared to single-data type subtypes.

**Fig 5.**
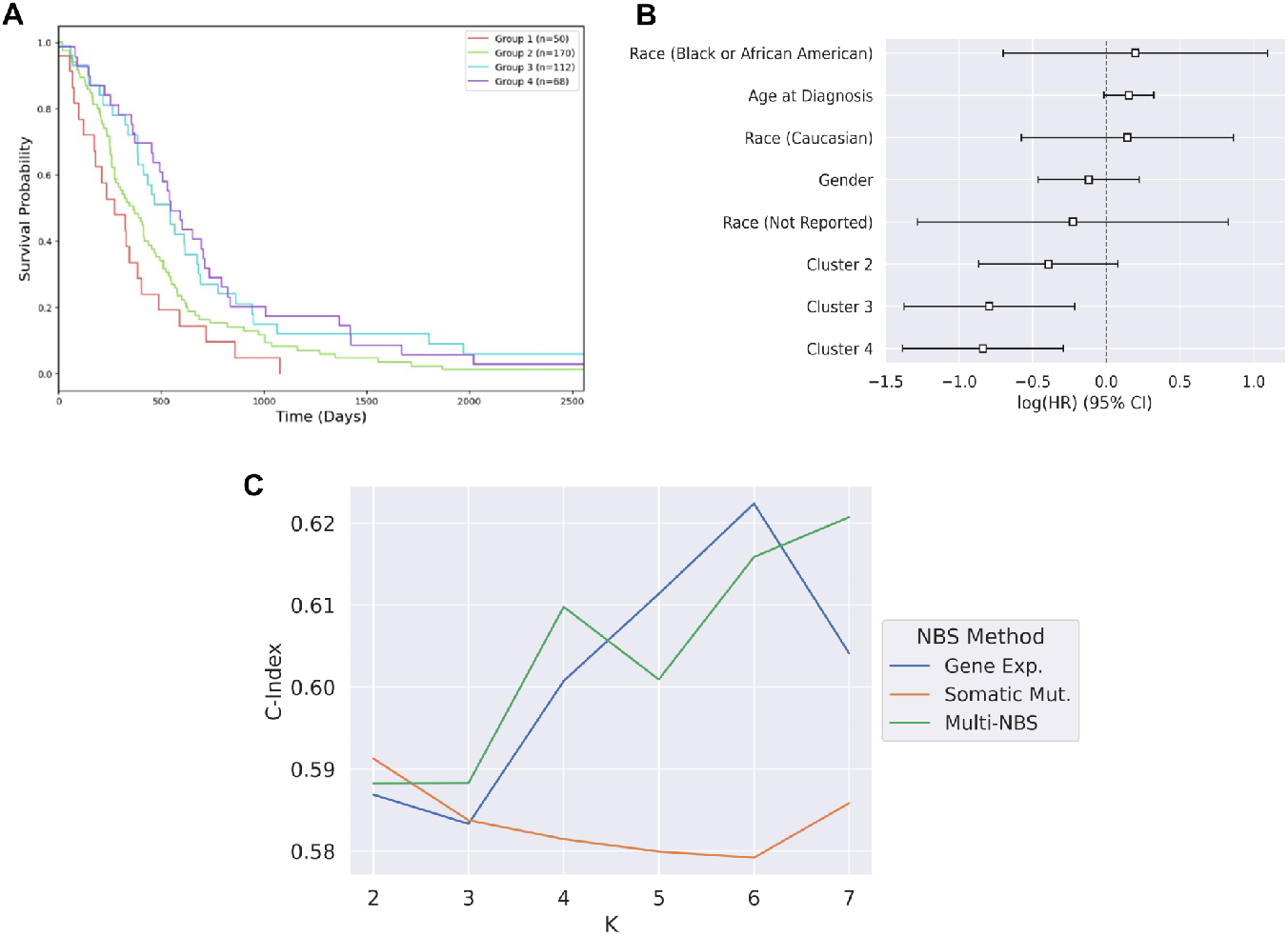
Bladder cancer survival analysis. Bladder cancer survival analysis indicate that integrated subtypes are significantly informative of survival. (A) Survival curves indicating the probability of survival for subtypes generated by Multi-NBS (log-rank test, *p* = 2 × 10^−3^). (B) Cox regression log hazard ratios for *K* = 4 integrated subtypes. Concordance index of different clusters for bladder cancer subtypes.

### Association with TCGA database subtypes

To further investigate the biological significance, we applied *χ*^2^ association tests with subtypes derived from other data types in the TCGA, including copy-number variation (CNV), methylation, mRNA expression, microRNA expression and protein profiles [23–25]. These TCGA subtypes provided by the TCGA database are determined through analyzing various molecular and histological factors [26–28]. For more information, refer to the Methods section.

For ovarian cancer, we see none of the generated clusters showed a significant association with TCGA subtypes in Fig 6A. This indicates that the single-data type and Multi-NBS subtypes identified are independent of TCGA subtypes. Cluster similarity analysis between Multi-NBS subtypes and TCGA subtypes (AMI = −0.0013) also supports Multi-NBS subtype assignments being independent of TCGA subtypes. For bladder cancer, however, we found both TCGA subtypes and Multi-NBS subtypes are significantly associated with survival, though Multi-NBS is significantly associated with TCGA subtypes more consistently than single-data types with log-rank test, *p* = 1.16 *×* 10^−2^ and *p* = 4 *×* 10^−4^, respectively (Fig 6B, S4 Fig). For uterine cancer, we see Multi-NBS subtypes exhibited stronger and more consistent association with TCGA subtypes than single-data type clusters (Fig 6C). We also note that NBS subtypes produced from only somatic mutation profiles are also significantly associated with TCGA subtypes, which is in line with previous findings [13]. Overall, our analysis has revealed that integrated profile-generated subtypes through NBS are potentially more capable than single-data type NBS subtypes of forming stronger association with TCGA subtypes for bladder cancer and uterine cancer.

**Fig 6.**
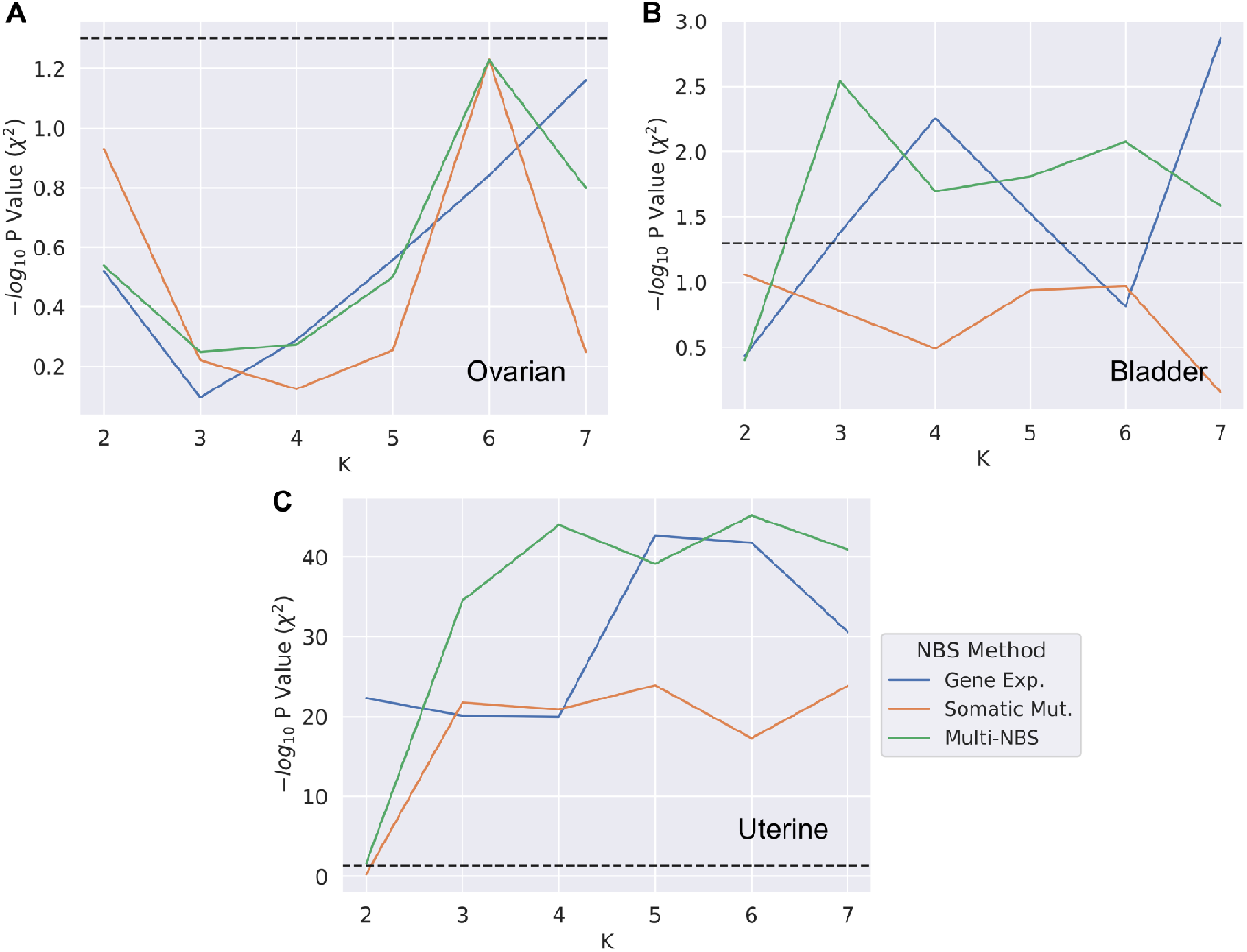
Association with TCGA subtypes. Association tests with TCGA provided subtypes across various cancers. The dotted line represents *p* = 0.05. Refer to the legend in subfigure (C) for all subfigures. (A) Generated ovarian cancer subtypes association with TCGA ovarian cancer subtypes. (B) Generated bladder cancer subtypes association with TCGA bladder cancer subtypes. (C) Generated uterine cancer subtypes association with TCGA uterine cancer subtypes.

### High-scoring genes

Next, we sought to identify genes with high scores across subtypes generated from Multi-NBS using network smoothed integrated profiles. Since the network values for integrated profile networks are a combination of gene expression and somatic mutation data, we refer to these scores as hybrid scores. We identified the genes with the highest average hybrid score per individual for each subtype. We then compared these findings to high-scoring genes across NBS subtypes generated from only somatic mutation profiles using network smoothed somatic mutation profiles. Again, we identified the genes with the highest average mutation score per individual for each subtype. We limited our investigation to the top 10 highest scoring genes (see Methods for further information on high-scoring genes identification). Across all three cancers, we found that integrated profiles identified more high-scoring genes shared by all subtypes than somatic mutation profiles. We see there were also genes with high scores unique to a subtype or a subset of subtypes.

Here we investigate the genes shared across all subtypes in integrated profiles that were not identified in somatic mutation only profiles to assess their impact on tumour initiation and progression. In ovarian cancer, we found integrated profiles identified BBC3 and UBC as two genes that had high scores across all Multi-NBS subtypes. We see somatic mutation only subtypes only identified NFKBIZ as a high-scoring gene shared by 3 of the 4 somatic mutation NBS subtypes. We found the BBC3 gene had the highest score across all 4 Multi-NBS subtypes. BBC3, or p53 upregulated modulator of apoptosis, whose expression is modulated by tumor suppressor p53. BBC3 has been identified as a strong marker for indicators of cancer [29]. UBC, which has a role in maintaining ubiquitin homeostasis under stress, also has a high score across all subtypes. Current research indicates that UBC is upregulated in cancers and plays a role in the increased proliferation rate of cancer cells and their ability to overcome cellular stresses introduced by cancer treatments [30]. We list the high-scoring genes for integrated NBS subtypes and somatic mutation NBS subtypes for ovarian cancer in S1 Table and S2 Table, respectively.

In bladder cancer, we observe integrated profiles identified ABCC1, LEPR, LTB4R, and IL1RAP as high-scoring genes across all 4 Multi-NBS subtypes while somatic mutation profiles identified ABCC1 and IL1RAP as high-scoring genes present across all 4 NBS subtypes generated from somatic mutation profiles only (S3 Table and S4 Table). LEPR is understood to be involved in tumour proliferation and migration through regulating ERK1/ERK2 and JAK2/STAT3 expression by interacting with ANXA7 [31]. Recently, there has been literature surrounding the possibility of a variant of LEPR being involved in bladder cancer due to the role of leptin in obese individuals [32]. LTB4R, or leukotriene B4 receptors, are involved in cell proliferation, survival, and metastasis. Previous literature has identified LTB4R as a potential biomarker for tumour sensitivity [33].

In uterine cancer, we found integrated profiles identified IDO1, NCK2, EDAR, and DUSP8 as high-scoring genes across all 3 Multi-NBS subtypes. We observe somatic mutation profiles only identified IDO1 as a high-scoring gene shared by all 3 NBS subtypes generated from somatic mutation profiles. NCK2 is highly involved in pathways mediating cytoskeleton organization and cell proliferation with previous literature indicating NCK2 expression is linked to metastatic human melanoma tumours [34]. EDAR is part for the Tumour Necrosis Factor Receptor (TNFR) superfamily and is a death receptor. Research has identified EDAR as an oncogene for breast cancer, and could be a potential target for investigation in other cancers [35]. DUSP8 is involved in the mitogen-activated protein kinase (MAPK) signaling pathway and has been shown to be tumour progression and drug resistance in various cancers such as colorectal cancer, lung cancer, and breast cancer [36–38]. These previous findings indicate that DUSP8 could be a potential candidate for its involvement in uterine cancer. We list the high-scoring genes for integrated NBS subtypes and somatic mutation NBS subtypes for uterine cancer in S5 Table and S6 Table, respectively.

In summary, we found integrated NBS subtypes exhibit a potentially enhanced capability in identifying highly significant genes shared across all subtypes compared to somatic mutation NBS subtypes. The observed shared high-scoring genes are unique for each type of cancer, as evidenced by our analysis which revealed no overlap in these genes among the three cancer types studied. Moreover, the genes within the integrated networks have a documented role in promoting tumor initiation, progression, and drug resistance, as supported by our investigation of existing literature. This underscores the potential of integrated NBS to pinpoint cancer-specific genes warranting further investigation.

## Discussion

In this study, we proposed a method that integrates RNA-seq and somatic mutation profiles before undergoing the NBS process to stratify tumors in an unsupervised manner. Through subsequent analysis, we identified the integrated clusters form comparably separated clusters compared to single-data type clusters. We found this integrated approach is also more effective compared to single-data type NBS in producing subtypes associated with survival, even after accounting for clinical covariates. We see this integrated approach also yielded subtypes more significantly associated with TCGA provided subtypes. We also found this integrated approach identified key genes with high mutation scores that were specific to a subtype but also other genes that were highly prevalent across all subtypes.

The reason Multi-NBS performs well could be largely due to the more holistic nature of integrating somatic mutation and RNA-seq profiles. Somatic mutation profiles are a relative comparison of healthy and tumor tissue, while RNA-seq profiles are an absolute measure of the cell state. Somatic mutations are useful because they contain casual genetic signals driving tumor progression and are differential measures between cancer tumors and normal tissue, which is more capable of capturing tumor-specific alterations. However, the tumor phenotype is often variable even when causal genes are identified, and RNA-seq profiles potentially offer to bridge this gap between phenotype and causal genes by better capturing highly expressed genes. Notably, we see integrated profiles were able to identify more genes that had high network scores across all subtypes within a cancer type. Thus, integrated profiles provide a more comprehensive view of the molecular processes underlying tumor initiation and progression, resulting in the improved stratification we observed.

It is crucial to acknowledge that the choice of *β* values may not remain consistent across all cancers, revealing a potential vulnerability of the method. This variability underscores the importance of tailoring the integration process to the unique characteristics of each cancer type for subtype analysis. In our study, integrating somatic mutation and gene expression profiles involved tuning the hyperparameter *β* to balance these two data types which led to Multi-NBS subtypes exhibiting improved biological and clinical relevance. However, determining optimal *β* values proved challenging and varied across cancers. For ovarian, bladder, and uterine cancers, we utilized tuned *β* values of 0.8, 0.3, and 0.1, respectively, guided by survival analysis metrics or TCGA subtype association metrics. The fluctuation of *β* across cancers emphasizes the need for careful consideration and adaptation when implementing Multi-NBS for cancer subtyping to suit the specific biological nuances of each cancer type.

There are still many methods to integrate different layers of genetic data within the NBS framework. Multiple additional layers of genetic data such as copy-number variation and protein expression data could be integrated similarly to identify clinically and biologically relevant subtypes. Another avenue of exploration is the integration of different genetic data that is beyond a linear combination of genetic profiles. This could include using protein-protein interaction, metabolic, or signaling networks in combination with genetic profiles. Exploration of NBS-generated subtypes through different genetic data integration methods could lead to more clinically and biologically informative subtypes. Investigating these integrated NBS approaches across a wide variety of cancers can lead to a stronger understanding of tumor initiation and progression.

## Acknowledgments

The research reported in this paper was supported by the NSF Quantitative Biology REU Site at Northwestern (DMS-2150134); Northwestern Research Training Grant in Quantitative Biological Modeling, (NSF DMS-1547394); NSF-Simons Center for Quantitative Biology (Simons Foundation/SFARI 597491-RWC and NSF DMS-1764421). The authors gratefully acknowledge the support provided by these grants.

## Supporting information

**S1 Fig.**
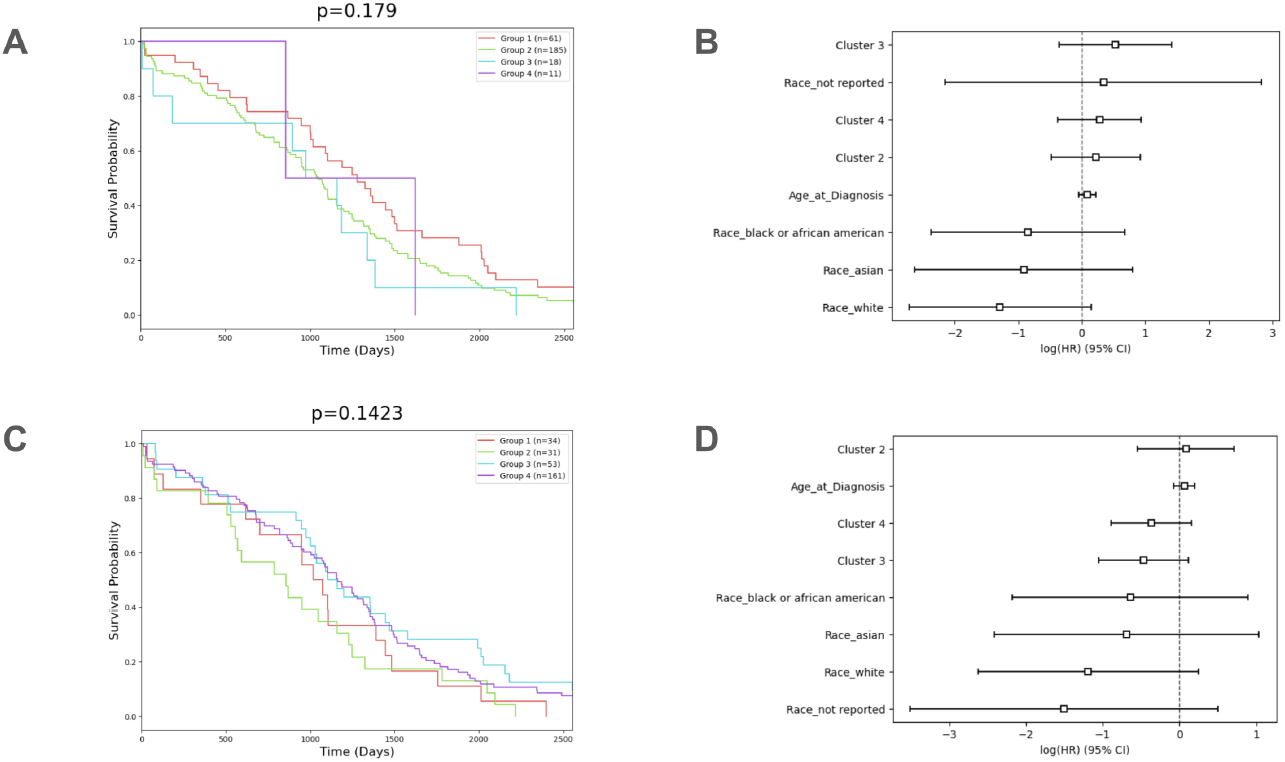
Ovarian cancer single-data type survival analysis. (A, C) Ovarian cancer single-data type survival curves for *K* = 4 with the log-rank test p-value above each survival curve. Single-data type subtypes are not significantly associated with survival while integrated profile generated subtypes are significantly associated with survival as shown in the paper. (B, D) Ovarian cancer log hazard ratios for single-data type generated clusters. The cluster coefficients are not significantly nonzero indicating the cluster assignment is not informative of higher or lower survival than subtype 1.

**S2 Fig.**
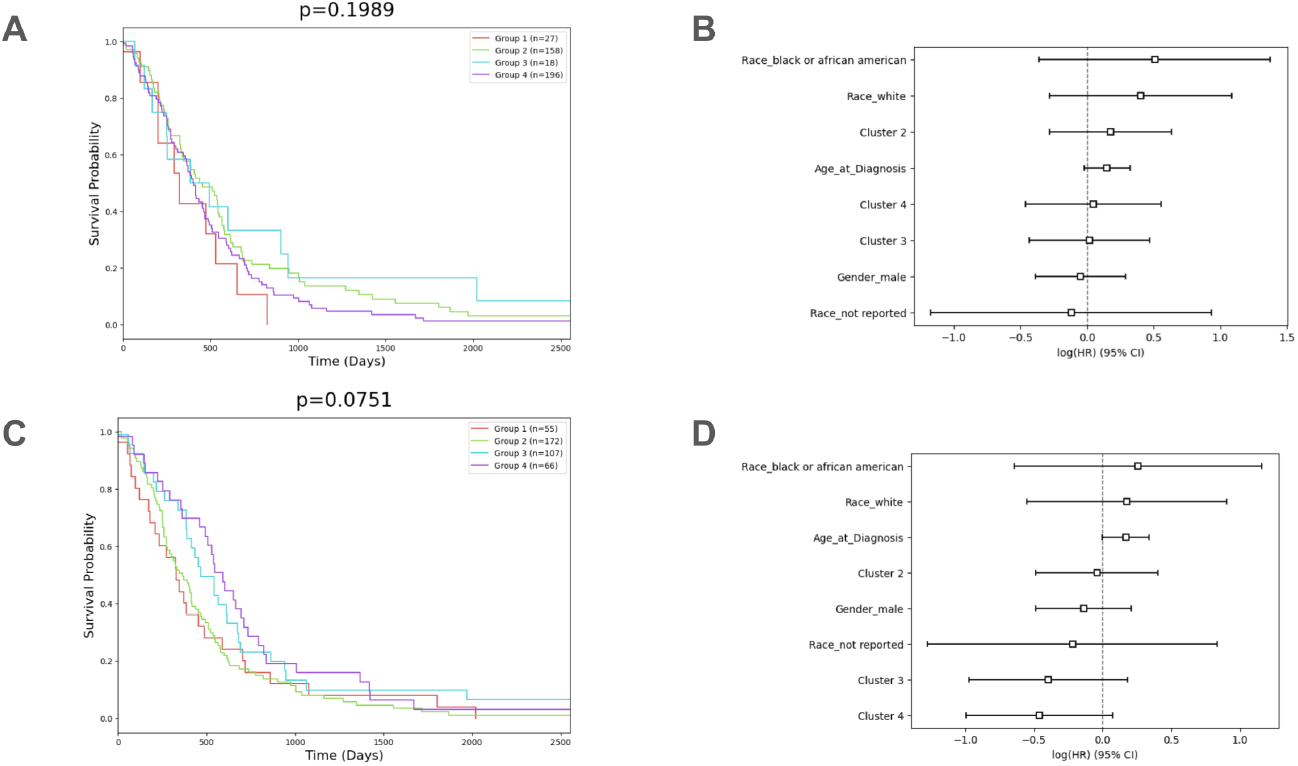
Bladder cancer single-data type survival analysis. (A, C) Bladder cancer single-data type survival curves for *K* = 3 with the log-rank test p-value above each survival curve. Single-data type subtypes are not significantly associated with survival while integrated profile generated subtypes are significantly associated with survival as shown in the paper. (B, D) Bladder cancer log hazard ratios for single-data type generated clusters. The cluster coefficients are not significantly nonzero indicating the cluster assignment is not informative of higher or lower survival than subtype 1.

**S3 Fig.**
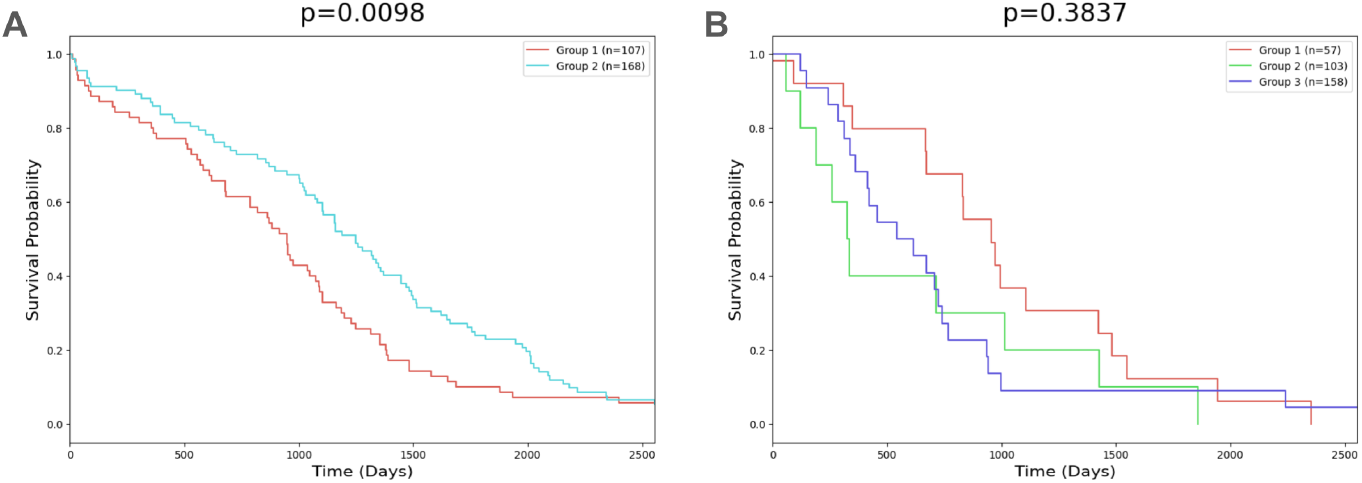
Somatic mutation generated subtypes. NBS subtypes using only somatic mutation profiles for ovarian cancer (*K* = 2) and uterine cancer (*K* = 3) with the associated survival log-rank test p-value.

**S4 Fig.**
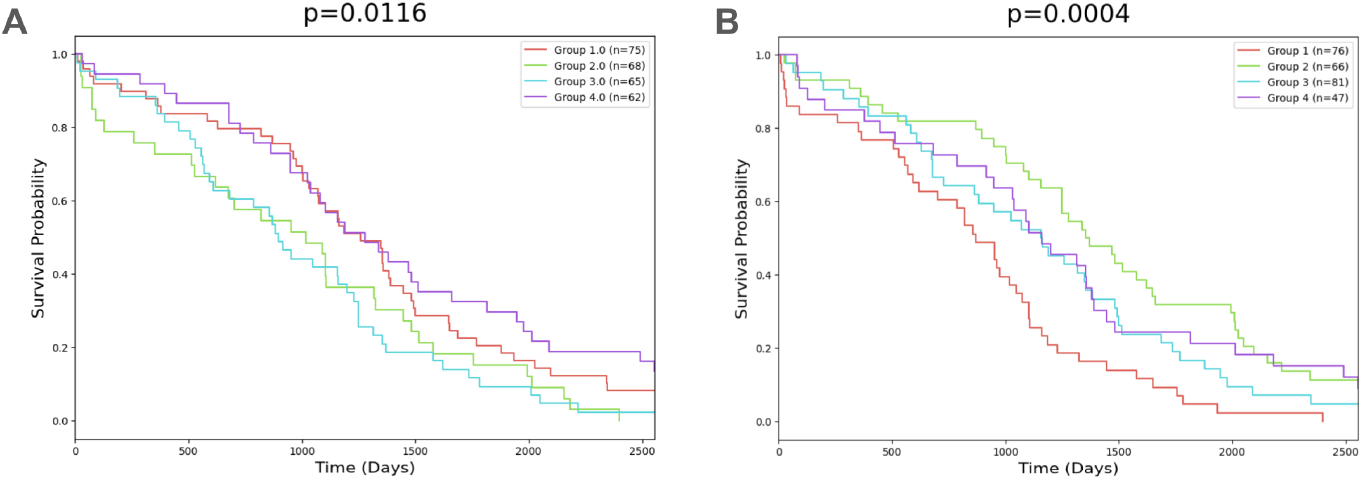
TCGA subtypes and Multi-NBS subtype survival curves. Survival curves from ovarian cancer TCGA subtypes and Multi-NBS generated subtypes (*K* = 4). Both are significantly associated with survival through log-rank test.

**S1 Table.** Integrated ovarian subtypes hybrid scores. The genes with the highest mean hybrid score across the networks of the patients in ovarian cancer subtypes using integrated profiles. BBC3 and UBC are relevant high-scoring genes across all 4 subtypes.

| Rank | Subtype 1 |  | Subtype 2 |  | Subtype 3 |  | Subtype 4 |  |
| --- | --- | --- | --- | --- | --- | --- | --- | --- |
|  | Gene | Score | Gene | Score | Gene | Score | Gene | Score |
| 1 | BBC3 | 0.159 | BBC3 | 0.146 | BBC3 | 0.166 | BBC3 | 0.157 |
| 2 | UBC | 0.127 | MAPK11 | 0.072 | UBC | 0.110 | UBC | 0.121 |
| 3 | CCNB1 | 0.099 | UBC | 0.071 | JAK3 | 0.097 | RB1 | 0.097 |
| 4 | YWHAB | 0.099 | ORC5 | 0.068 | HRAS | 0.096 | KRAS | 0.091 |
| 5 | DHX9 | 0.098 | NOTCH3 | 0.068 | GSK3A | 0.093 | HRAS | 0.091 |
| 6 | HSP90AB1 | 0.097 | EIF3E | 0.066 | IL6 | 0.091 | LIF | 0.091 |
| 7 | TNF | 0.096 | PPP3R1 | 0.066 | RB1 | 0.091 | JAK3 | 0.091 |
| 8 | RB1 | 0.095 | NAP1L1 | 0.065 | SMC3 | 0.090 | MYL7 | 0.089 |
| 9 | TERT | 0.095 | IFNG | 0.064 | STAT3 | 0.090 | A2M | 0.088 |
| 10 | HRAS | 0.092 | PPP2R5B | 0.064 | KRAS | 0.090 | FN1 | 0.088 |

**S2 Table.**
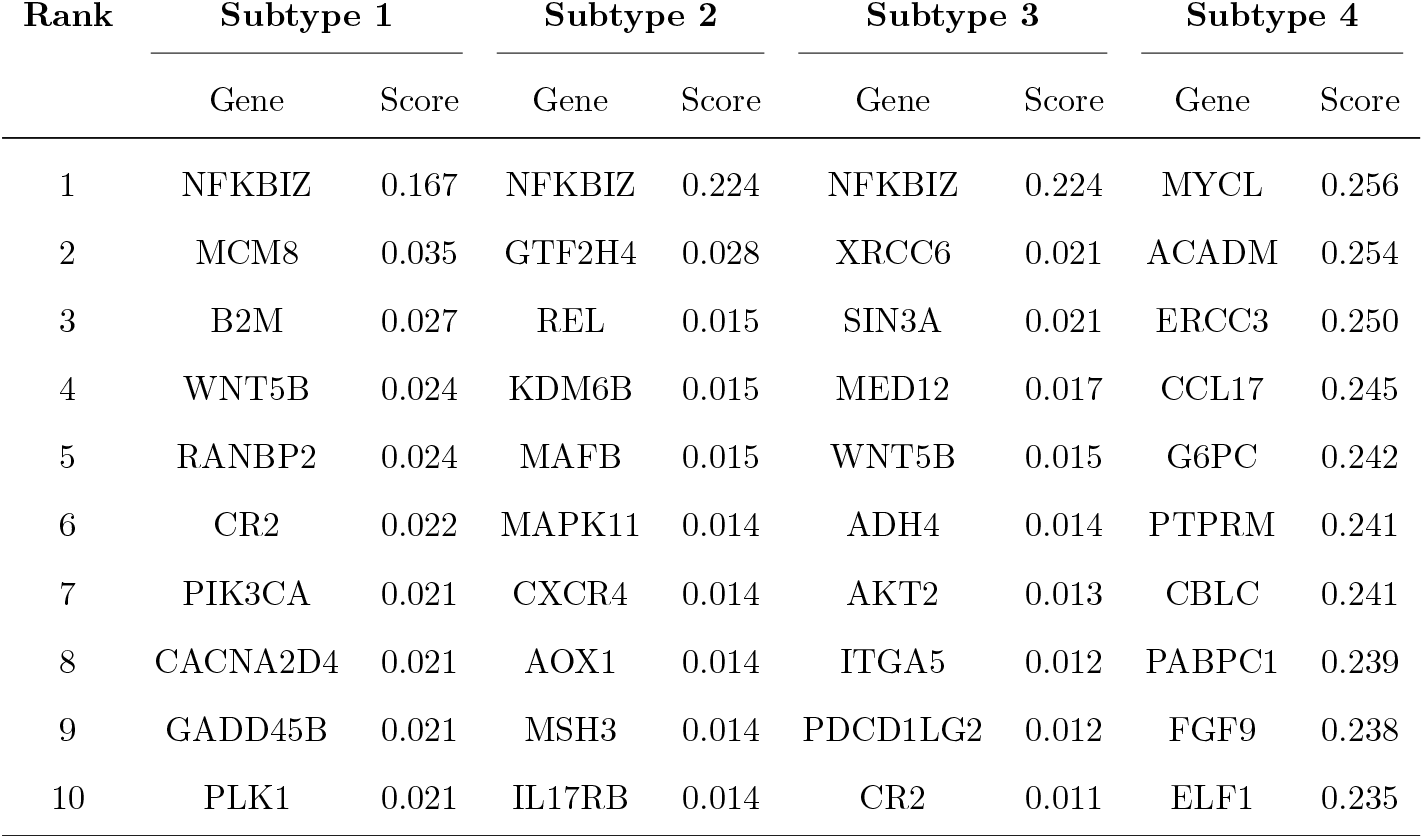
Somatic mutation ovarian subtypes mutation scores. The genes with the highest mean mutation score across the networks of the patients in ovarian cancer subtypes using only somatic mutation profiles profiles. NFKBIZ is a relevant high-scoring genes across 3 of the 4 subtypes.

**S3 Table.** Integrated bladder subtypes hybrid scores. The genes with the highest mean hybrid score across the networks of the patients in bladder cancer subtypes using integrated profiles. ABCC1, LEPR, LTB4R, and IL1RAP are relevant high-scoring genes across all 4 subtypes.

| Rank | Subtype 1 |  | Subtype 2 |  | Subtype 3 |  | Subtype 4 |  |
| --- | --- | --- | --- | --- | --- | --- | --- | --- |
|  | Gene | Score | Gene | Score | Gene | Score | Gene | Score |
| 1 | ABCC1 | 0.155 | ABCC1 | 0.157 | LEPR | 0.090 | LTB4R | 0.087 |
| 2 | CCNH | 0.105 | IL1RAP | 0.095 | ABCC1 | 0.077 | ABCC1 | 0.079 |
| 3 | IL1RAP | 0.069 | LEPR | 0.076 | LTB4R | 0.067 | IL1RAP | 0.060 |
| 4 | AFAP1L2 | 0.055 | CCNH | 0.075 | IL1RAP | 0.060 | AFAP1L2 | 0.049 |
| 5 | IBSP | 0.054 | LTB4R | 0.066 | KLK3 | 0.057 | KLK3 | 0.044 |
| 6 | PICALM | 0.048 | AFAP1L2 | 0.059 | KIR2DS5 | 0.054 | LEPR | 0.043 |
| 7 | THBS3 | 0.045 | PICALM | 0.050 | ACAD8 | 0.050 | NFE2L2 | 0.041 |
| 8 | TICAM2 | 0.045 | HLA-E | 0.047 | HLA-E | 0.046 | RBM15 | 0.038 |
| 9 | LTB4R | 0.045 | KIR2DS5 | 0.046 | SF3B1 | 0.045 | PICALM | 0.038 |
| 10 | LEPR | 0.042 | CXCL1 | 0.046 | NTHL1 | 0.042 | ACAD8 | 0.036 |

**S4 Table.** Somatic mutation bladder subtypes mutation scores. The genes with the highest mean mutation score across the networks of the patients in bladder cancer subtypes using only somatic mutation profiles profiles.. ABCC1 and IL1RAP are relevant high-scoring genes across all 4 subtypes. LEPR is a high scoring gene across 3 of the 4 subtypes.

| Rank | Subtype 1 |  | Subtype 2 |  | Subtype 3 |  | Subtype 4 |  |
| --- | --- | --- | --- | --- | --- | --- | --- | --- |
|  | Gene | Score | Gene | Score | Gene | Score | Gene | Score |
| 1 | TPO | 0.112 | LEPR | 0.112 | ITGA2 | 0.112 | ABCC1 | 0.143 |
| 2 | NTHL1 | 0.087 | ABCC1 | 0.110 | ABCC1 | 0.108 | LTB4R | 0.112 |
| 3 | FGF18 | 0.049 | IL1RAP | 0.068 | CCNH | 0.091 | IL1RAP | 0.092 |
| 4 | LEPR | 0.048 | ACAD8 | 0.063 | KLK3 | 0.076 | CCNH | 0.067 |
| 5 | SF3B1 | 0.047 | KIR2DS5 | 0.058 | LTB4R | 0.074 | AFAP1L2 | 0.053 |
| 6 | ABCC1 | 0.046 | AFAP1L2 | 0.057 | MAPK12 | 0.056 | HLA-E | 0.048 |
| 7 | MMP1 | 0.040 | KLK3 | 0.046 | NTN4 | 0.055 | CXCL1 | 0.048 |
| 8 | PICALM | 0.037 | PICALM-E | 0.043 | RFC5 | 0.047 | PICALM | 0.043 |
| 9 | TYROBP | 0.033 | THBS3 | 0.042 | IL1RAP | 0.047 | SF3B1 | 0.042 |
| 10 | IL1RAP | 0.032 | HLA-E | 0.040 | NTHL1 | 0.046 | LEPR | 0.041 |

**S5 Table.**
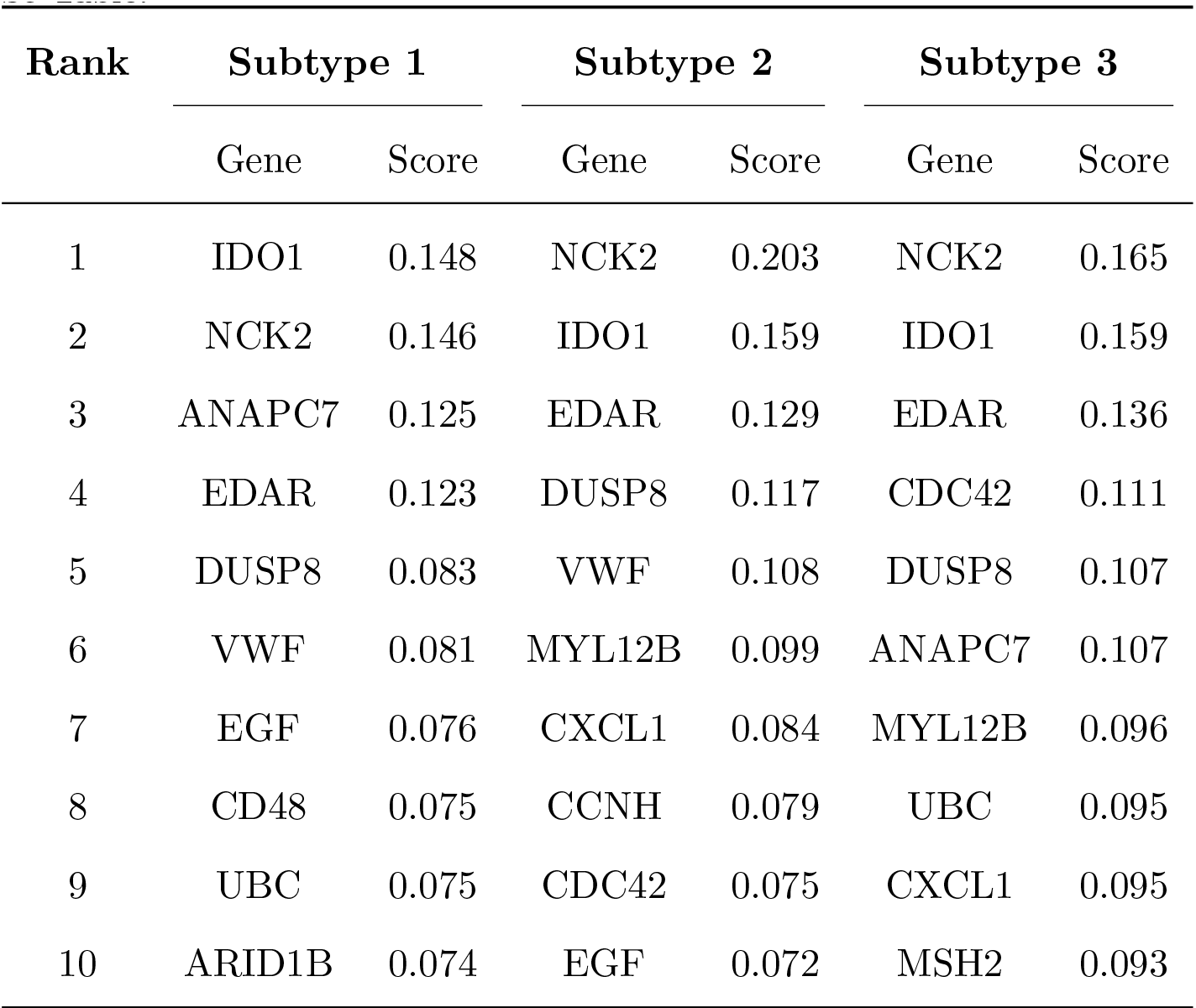
Integrated uterine subtypes hybrid scores. The genes with the highest mean hybrid score across the networks of the patients in uterine cancer subtypes using integrated profiles. IDO1, NCK2, EDAR, and DUSP8 are relevant high-scoring genes across all 3 subtypes.

**S6 Table.** Somatic mutation uterine subtypes mutation scores. The genes with the highest mean mutation score across the networks of the patients in uterine cancer subtypes using only somatic mutation profiles. IDO1 is a relevant high-scoring gene across all 3 subtypes. NCK2, EDAR, and DUSP8 are high-scoring genes in 2 out of the 3 subtypes.

| Rank | Subtype 1 |  | Subtype 2 |  | Subtype 3 |  |
| --- | --- | --- | --- | --- | --- | --- |
|  | Gene | Score | Gene | Score | Gene | Score |
| 1 | ANAPC7 | 0.247 | NCK2 | 0.187 | NCK2 | 0.219 |
| 2 | IDO1 | 0.112 | IDO1 | 0.140 | EDAR | 0.191 |
| 3 | TNFRSF13C | 0.083 | MYL12B | 0.119 | IDO1 | 0.183 |
| 4 | ARID1B | 0.058 | EDAR | 0.110 | CDC42 | 0.179 |
| 5 | CCL8 | 0.037 | DUSP8 | 0.088 | EGF | 0.164 |
| 6 | DUSP8 | 0.036 | VWF | 0.066 | UBC | 0.162 |
| 7 | IL17RA | 0.031 | PAX8 | 0.034 | MSH2 | 0.161 |
| 8 | CCR9 | 0.030 | CCNH | 0.032 | CXCL1 | 0.161 |
| 9 | CNBP | 0.022 | CCL8 | 0.026 | YWHAQ | 0.147 |
| 10 | LRP6 | 0.016 | CXCL16 | 0.025 | CHUK | 0.146 |

